# New auranofin analogs with antibacterial properties against *Burkholderia* clinical isolates

**DOI:** 10.1101/2021.09.10.459877

**Authors:** Dustin Maydaniuk, Bin Wu, Dang Truong, Sajani H. Liyanage, Andrew M. Hogan, Zhong Ling Yap, Mingdi Yan, Silvia T. Cardona

## Abstract

Bacteria of the genus *Burkholderia* include pathogenic *Burkholderia mallei, Burkholderia pseudomallei* and the *Burkholderia cepacia* complex (Bcc). These Gram-negative pathogens have intrinsic drug resistance, which makes treatment of infections difficult. Bcc affects individuals with cystic fibrosis (CF) and the species *B. cenocepacia* is associated with one of the worst clinical outcomes. Following the repurposing of auranofin as an antibacterial against Gram-positive bacteria, we previously synthetized auranofin analogs with activity against Gram-negatives. In this work, we show that two auranofin analogs, MS-40S and MS-40, have antibiotic activity against *Burkholderia* clinical isolates. The compounds are bactericidal against *B. cenocepacia* and kill stationary-phase cells and persisters without selecting for multistep resistance. *Caenorhabditis elegans* and *Galleria mellonella* tolerated high concentrations of MS-40S and MS-40, demonstrating that these compounds have low toxicity in these model organisms. In summary, we show that MS-40 and MS-40S have the potential to be effective therapeutic options to treat *Burkholderia* infections.

## Introduction

Antibiotics are one of the greatest medical advances of the 20^th^ century, with their widespread discovery starting in the early 1900’s (1,2). However, antibiotic resistance is now a global crisis, responsible for approximately 700,000 deaths annually (2), with that number projecting to increase each year (3). The “golden era” of antibiotic discovery, which lasted approximately 20 years, led to the identification of vancomycin, methicillin, cephalosporins, and many other antibiotics (1,2). No new antibiotic from a new class has reached the clinics for many decades (4). When a new antibiotic is developed that doesn’t have a novel mechanism of action, resistance mechanisms are already present. Therefore, antimicrobials with novel mechanisms of action are needed to prevent this quickly generated resistance.

Bacteria of the genus *Burkholderia* (5) includes difficult-to-treat human pathogens such as the *Burkholderia cepacia* complex (Bcc), *Burkholderia mallei* and *Burkholderia pseudomallei* (6). Bcc is a group of more than 20 species that cause life threatening bacterial infections in cystic fibrosis (CF) patients (7). *Burkholderia cenocepacia* infections, in particular, have one of the worst clinical outcomes (5, 8), causing decreased lung function and *cepacia* syndrome, a sepsis with necrotizing pneumonia (8, 9). Additionally, CF patients infected with *B. cenocepacia* are often ineligible for lung transplants (10), a common life-saving procedure (8), because *B. cenocepacia* infections have a high risk of reoccurrence (11).

New antibiotics are urgently needed to combat *Burkholderia* infections. Promising emerging therapeutics are those with heavy metals (12), especially gold (13–17), which have been used in medicine since 2500 B.C.E. Recently, the gold-containing, anti-arthritis drug auranofin (18, 19) was found to be active against *Mycobacterium tuberculosis* and Gram-positive bacteria (20). Auranofin inhibited the function of the enzyme thioredoxin reductase, interrupting thiol-redox homeostasis (21,22). However, auranofin lacked significant activity against Gram-negative bacteria with minimum inhibitory concentrations (MICs) >16 mg/L (20).

We previously found that auranofin is active against *Helicobacter pylori* and synthesized sugar-modified analogs with improved antibiotic activity and reduced toxicity to mammalian cells (21). By varying the structures of the thiol and phosphine ligands on auranofin, we expanded the antibacterial activity of the auranofin analogs to the Gram-negative pathogens including *Klebsiella pneumoniae, Acinetobacter baumannii, Pseudomonas aeruginosa*, and *Escherichia coli* (22). In this work, we describe the characterization of two auranofin analogs, WB-19-HL4170 (MS-40S) and WB-19-HL4118 (MS-40), which show potential as antimicrobials against Bcc strains, *B. pseudomallei* and *B. mallei*. MS-40S and MS-40 are bactericidal against *B. cenocepacia*, kill persister cells, and do not select for multistep resistant mutations while maintaining low toxicity to eukaryotic systems.

## Results

### MIC of auranofin derivatives against a panel of *Burkholderia cepacia* complex species

We tested auranofin and ten auranofin analogs (Figure 1) against a panel of Bcc bacteria, comprising of clinical isolates from CF patients and strains from environmental sources. Auranofin (Figure 1, top left) and the analogues belonging to group one (Figure 1, top right) were largely inactive against members of the Bcc (Table S1), with most of the MIC’s being 128 μg/mL or higher. Group one analogs have modifications of the thioglucose ligands, with or without additional replacing of trimethylphosphine (-PMe_3_) to triethylphosphine (-PEt_3_) coordinated bonding to the gold atom. Auranofin, however, showed high activity against *B. mallei* with MICs ranging from 0.25 – 1 μg/mL. In *B. pseudomallei*, auranofin had a MIC of 64 μg/mL. Auranofin and the group one derivatives in this study, where shown to have diverse activity in other Gram-negative and Gram-positive bacteria, but they were inactive or had low activity in *P. aeruginosa, Enterobacter cloacae*, and *K. pneumoniae* (22).

**Figure 1.**
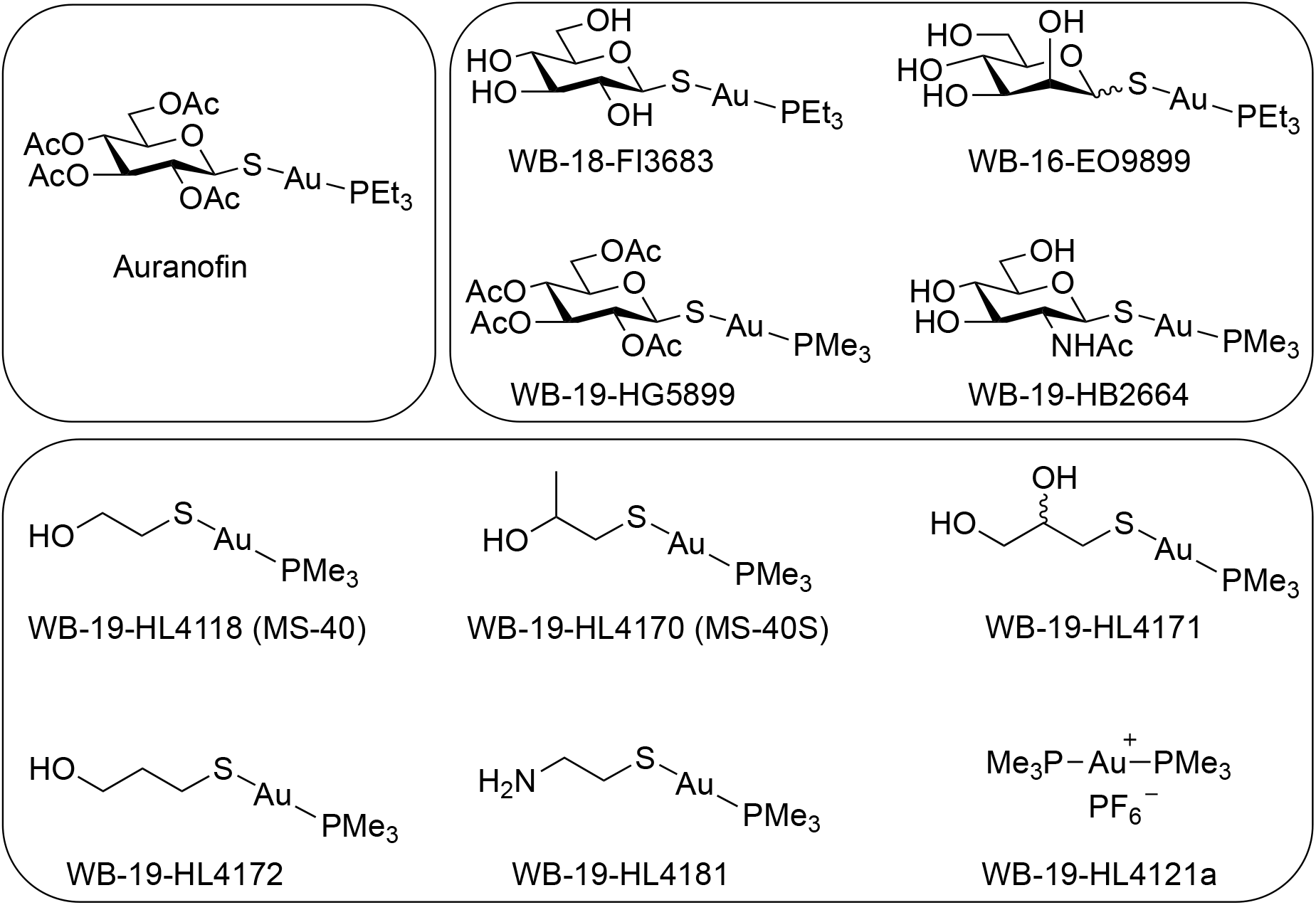
Chemical structure of auranofin and auranofin analogs. (Top left) Auranofin. (Top right) Group one auranofin analogs containing modifications of the thioglucose and replacement of triethylphosphine (P(CH_2_CH_3_)_3_; PEt_3_) with trimethylphosphine (P(CH_3_)_3_; PMe_3_). (Bottom) Group two auranofin analogs.

Group two (Figure 1, bottom) includes the analog WB-19-HL4118 (MS-40) which had shown a low MIC in Gram-negative and Gram-positive bacteria, such as in *A. baumannii, E. cloacae, E. coli*, and *S. aureus* (22).Therefore, we synthetized additional analogs with similar structures to MS-40. Group two analogs showed lower MICs against members of the Bcc (Table 1) than Group one (Table S1). Remarkably, MS-40 and WB-19-HL4170 (MS-40S) showed the strongest activity (Table 1). These two compounds, also have high activity in *B. mallei* and *B. pseudomallei* strains as well. The synthesis of MS-40S and MS-40 is shown in Scheme 1. The remaining derivatives from this group have moderate MICs, ranging from 8 to 64 μg/mL, with only a few being 128 μg/mL or higher. The structures of the group two derivatives have substitution of thioglucose ligands with mercaptoethanol (HOCH_2_CH_2_SH) or mercaptoethanol modification, suggesting that the thioglucose was unable to permeate into most of the Bcc bacterial cell.

**Table 1.**
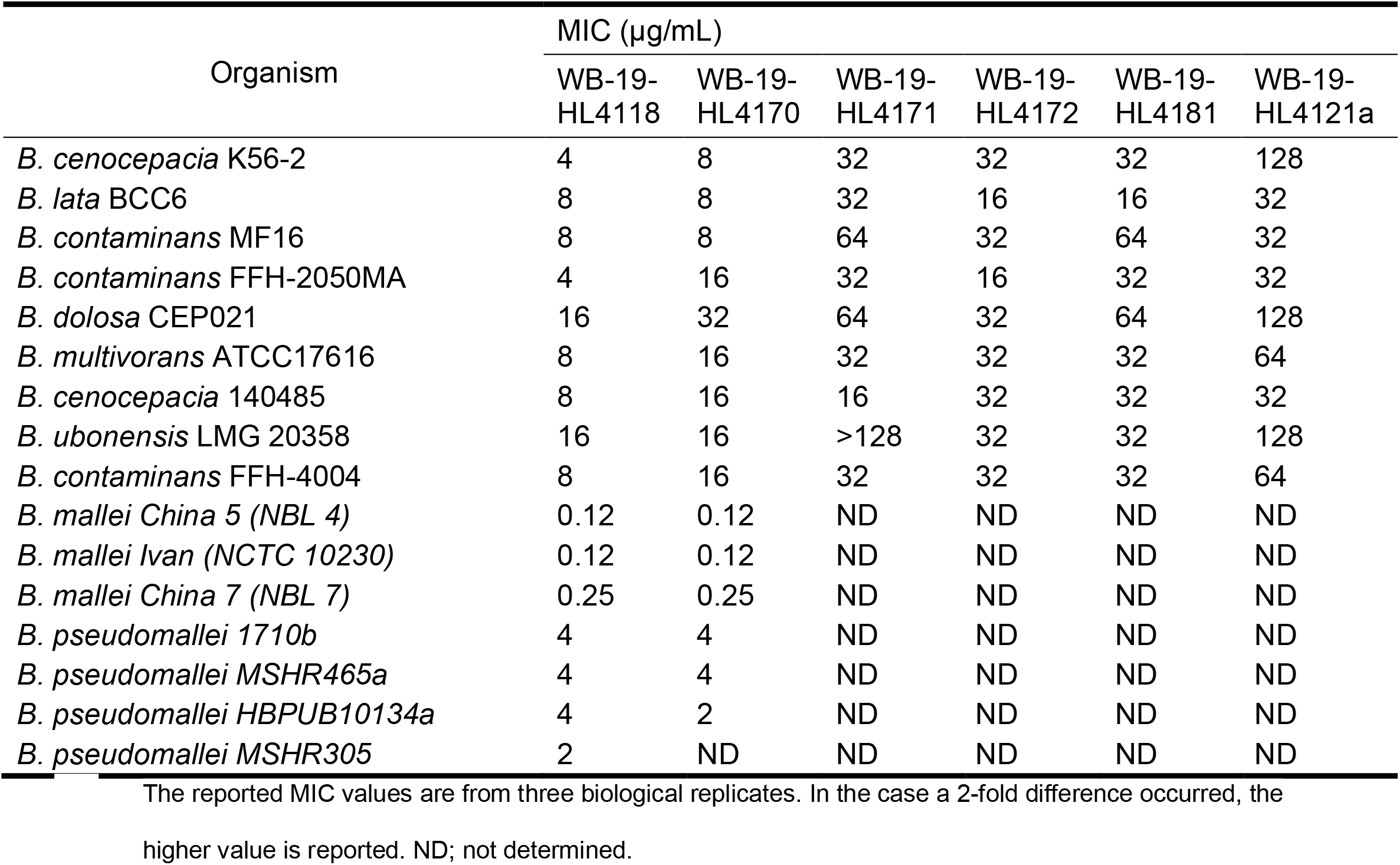
Minimum inhibitory concentrations (MICs) of group two auranofin derivatives against *Burkholderia cepacia* complex (Bcc) bacteria, *B. mallei* and *B. pseudomallei*.

| Organism | MIC (µg/mL) |  |  |  |  |  |
| --- | --- | --- | --- | --- | --- | --- |
|  | WB-19-HL4118 | WB-19-HL4170 | WB-19-HL4171 | WB-19-HL4172 | WB-19-HL4181 | WB-19-HL4121a |
| *B. cenocepacia* K56-2 | 4 | 8 | 32 | 32 | 32 | 128 |
| *B. lata* BCC6 | 8 | 8 | 32 | 16 | 16 | 32 |
| *B. contaminans* MF16 | 8 | 8 | 64 | 32 | 64 | 32 |
| *B. contaminans* FFH-2050MA | 4 | 16 | 32 | 16 | 32 | 32 |
| *B. dolosa* CEP021 | 16 | 32 | 64 | 32 | 64 | 128 |
| *B. multivorans* ATCC17616 | 8 | 16 | 32 | 32 | 32 | 64 |
| *B. cenocepacia* 140485 | 8 | 16 | 16 | 32 | 32 | 32 |
| *B. ubonensis* LMG 20358 | 16 | 16 | >128 | 32 | 32 | 128 |
| *B. contaminans* FFH-4004 | 8 | 16 | 32 | 32 | 32 | 64 |
| *B. mallei* China 5 (NBL 4) | 0.12 | 0.12 | ND | ND | ND | ND |
| *B. mallei* Ivan (NCTC 10230) | 0.12 | 0.12 | ND | ND | ND | ND |
| *B. mallei* China 7 (NBL 7) | 0.25 | 0.25 | ND | ND | ND | ND |
| *B. pseudomallei* 1710b | 4 | 4 | ND | ND | ND | ND |
| *B. pseudomallei* MSHR465a | 4 | 4 | ND | ND | ND | ND |
| *B. pseudomallei* HBPUB10134a | 4 | 2 | ND | ND | ND | ND |
| *B. pseudomallei* MSHR305 | 2 | ND | ND | ND | ND | ND |

The reported MIC values are from three biological replicates. In the case a 2-fold difference occurred, the higher value is reported. ND; not determined.

**Scheme 1.**
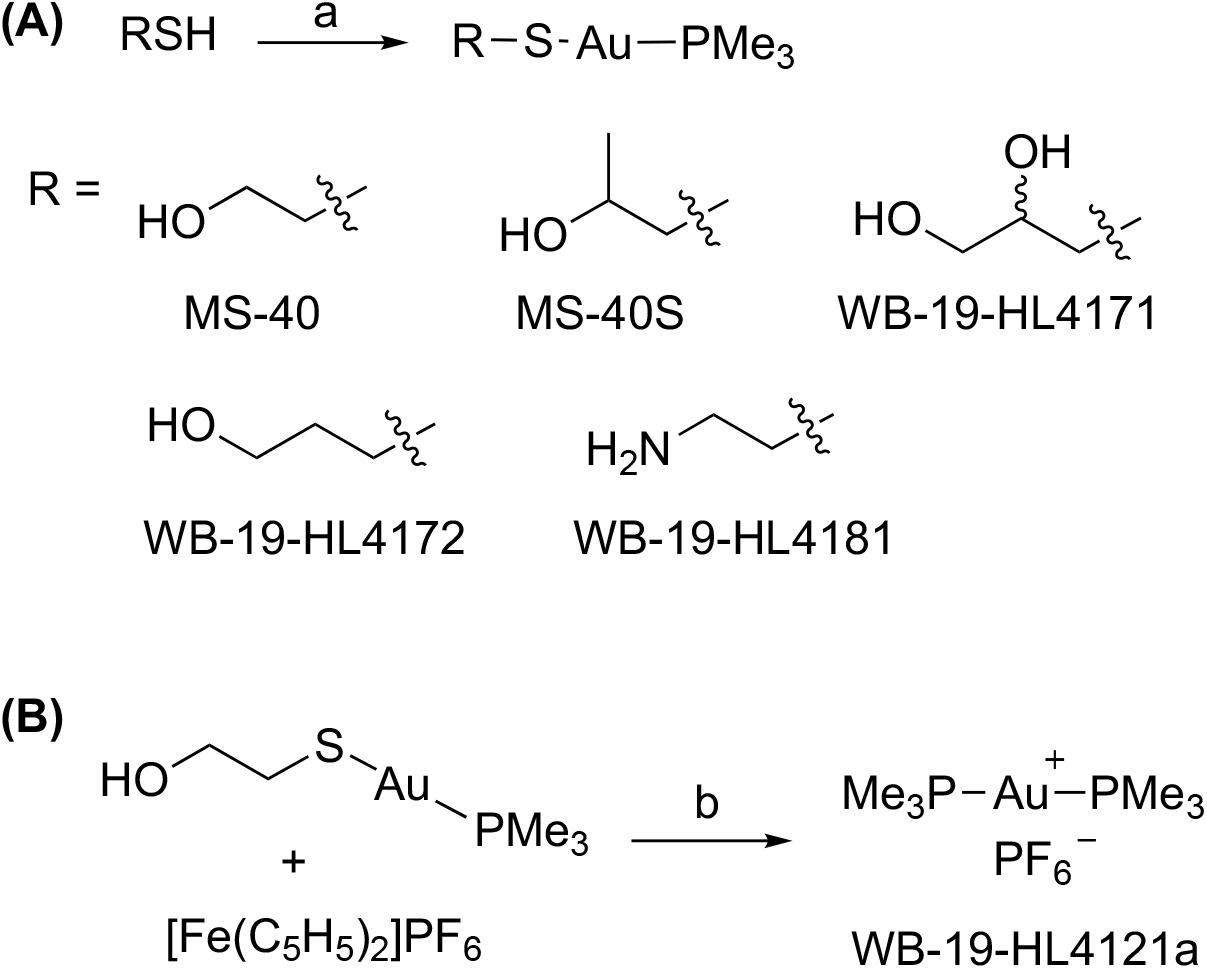
Synthesis of Group two compounds. Reagents and conditions: (a) NaOCH_3_ CH_3_OH, room temperature, 2 h; (b) dichloromethane, 0 °C, 23 h.

Next, we compared the MICs of MS-40S and MS-40 to common antibiotics used to treat CF patients infected with *Burkholderia spp*. Those include ceftazidime (23, 24), meropenem (23–26), doxycycline (27), and tobramycin (24–26). The combination therapy ceftazidime-avibactam is considered the last resort treatment for those infected with *Burkholderia* species(9); therefore, we determined the MIC of ceftazidime-avibactam and these four antibiotics against the Bcc panel (Table 2). The MICs of MS-40 and MS-40S are much lower than the antibiotic tobramycin and are similar to the other antibiotics, including the last resort combination treatment ceftazidime-avibactam. Additionally, doxycycline, against some isolates from the Bcc, had MIC values as low as 1 and 2 μg/mL. Taken together, the initial MIC testing shows MS-40S and MS-40 are comparable to antibiotics used currently in the clinic, having MIC’s lower than most and even have similar values as the combination therapy ceftazidime-avibactam.

**Table 2.** Minimum inhibitory concentrations (MICs) of MS-40S compared to clinical antibiotics used to treat CF patients.

| Organism | MIC (µg/mL) |  |  |  |  |  |  |
| --- | --- | --- | --- | --- | --- | --- | --- |
|  | MS-40 | MS-40S | MEM | TOB | CAZ | DOX | CZA<sup>a</sup> |
| *B. cenocepacia* K56-2 | 4 | 8 | 16 | 512 | 32 | 4 | 4 |
| *B. lata* BCC6 | 8 | 8 | 4 | 128 | 64 | 4 | 16 |
| *B. contaminans* MF16 | 8 | 8 | 8 | 32 | 32 | 16 | 8 |
| *B. contaminans* FFH-2050MA | 4 | 16 | 4 | 128 | 8 | 2 | 4 |
| *B. dolosa* CEP021 | 16 | 32 | 8 | 128 | 16 | 16 | 8 |
| *B. multivorans* ATCC17616 | 8 | 16 | 4 | 64 | 16 | 2 | 8 |
| *B. cenocepacia* 140485 | 8 | 16 | 16 | 128 | 8 | 1 | 8 |
| *B. ubonensis* LMG 20358 | 16 | 16 | 16 | 128 | 16 | 4 | 4 |
| *B. contaminans* FFH-4004 | 8 | 16 | 8 | 64 | 8 | 8 | 4 |

<sup>a</sup>8 µg/mL avibactam. The reported MIC values are from three biological replicates. In the case a 2-fold difference occurred, the higher value is reported. MEM, meropenem; TOB, tobramycin; CAZ, ceftazidime; DOX, doxycycline; CZA, ceftazidime-avibactam.

### MS-40S does not select for multistep resistant mutants

New antimicrobials are urgently needed because resistance to current antibiotics has arisen and spread to many bacteria (3, 28, 29). Ideally, resistance will occur slowly for new antimicrobials, or not at all. We therefore characterized the occurrence of resistance to MS-40S due to repeated exposure and continuous growth (30). Bacteria grown in the presence of subinhibitory concentrations of each compound (0.5× MIC) were subcultured and grown overnight in Luria-Bertani (LB) broth. These cultures were then used for the next MIC test, and this process was repeated for a total of 24 days (Figure 2, left). We performed this procedure for MS-40S, MS-40 and the antibiotics meropenem and doxycycline. These antibiotics were chosen because they are commonly used to treat cystic fibrosis patients infected with *Burkholderia* species (23–27), and they have different mechanisms of actions and resistance (3, 31).

**Figure 2.**
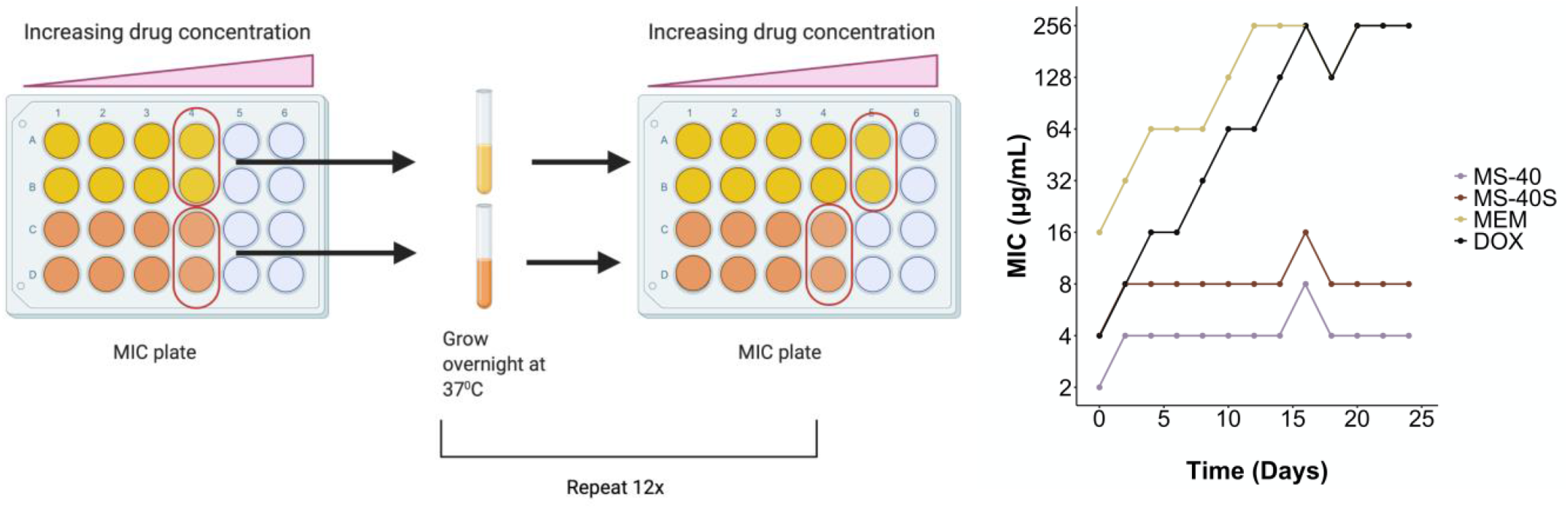
Resistance is not generated to MS-40S. Repeated exposure of a continuously grown culture to sub-lethal concentrations of the antimicrobials were achieved by determining the MIC of the compound. Then, for each compound, 30 μL of bacteria from the well with growth with the highest concentration of compound was grown overnight and used in the next MIC test. This was repeated over 24 days, with the MIC tested every second day. MEM, meropenem; DOX, doxycycline.

Figure 2 (right) shows that resistance against meropenem and doxycycline arose quickly. While their starting MICs values were 16 and 4 μg/mL, respectively, both MICs reached 256 μg/mL after 12-16 days. Remarkably, no apparent increase in their MICs was observed for MS-40S and MS-40, demonstrating a desired property as a potential therapeutic agent.

### MS-40S is bactericidal against both replicating and non-replicating cells

Antibiotics are classified either as bactericidal, if they kill cells and reduce the population by 99.99%, or bacteriostatic, if they prevent cell growth/division, but do not kill more than 99.99% of the population (31,32). It is common for antibiotics to only target actively dividing cells because their targets are involved in replication or other energy dependent processes (31), rendering them less effective when cells are not replicating or respiring (33). In time kill experiments, we found MS-40S to be bactericidal to both exponential (replicating) and stationary (non-replicating) phase cells, and the same was found for MS-40 (Figure 3; top 4 panels). Interestingly, MS-40S is more effective at killing stationary phase (Figure 3; middle right) than exponential phase cells (Figure 3; middle left), reducing the culture by approximately three log_10_ units and nine log_10_ units at 4× MIC in exponential and stationary phase, respectively. For comparison purposes, we show that doxycycline and ceftazidime-avibactam are both unable to kill cells in stationary phase (Figure 3; bottom right) and ceftazidime-avibactam is slow at killing exponential phase cells, regardless of the concentration (Figure 3; bottom left).

**Figure 3.**
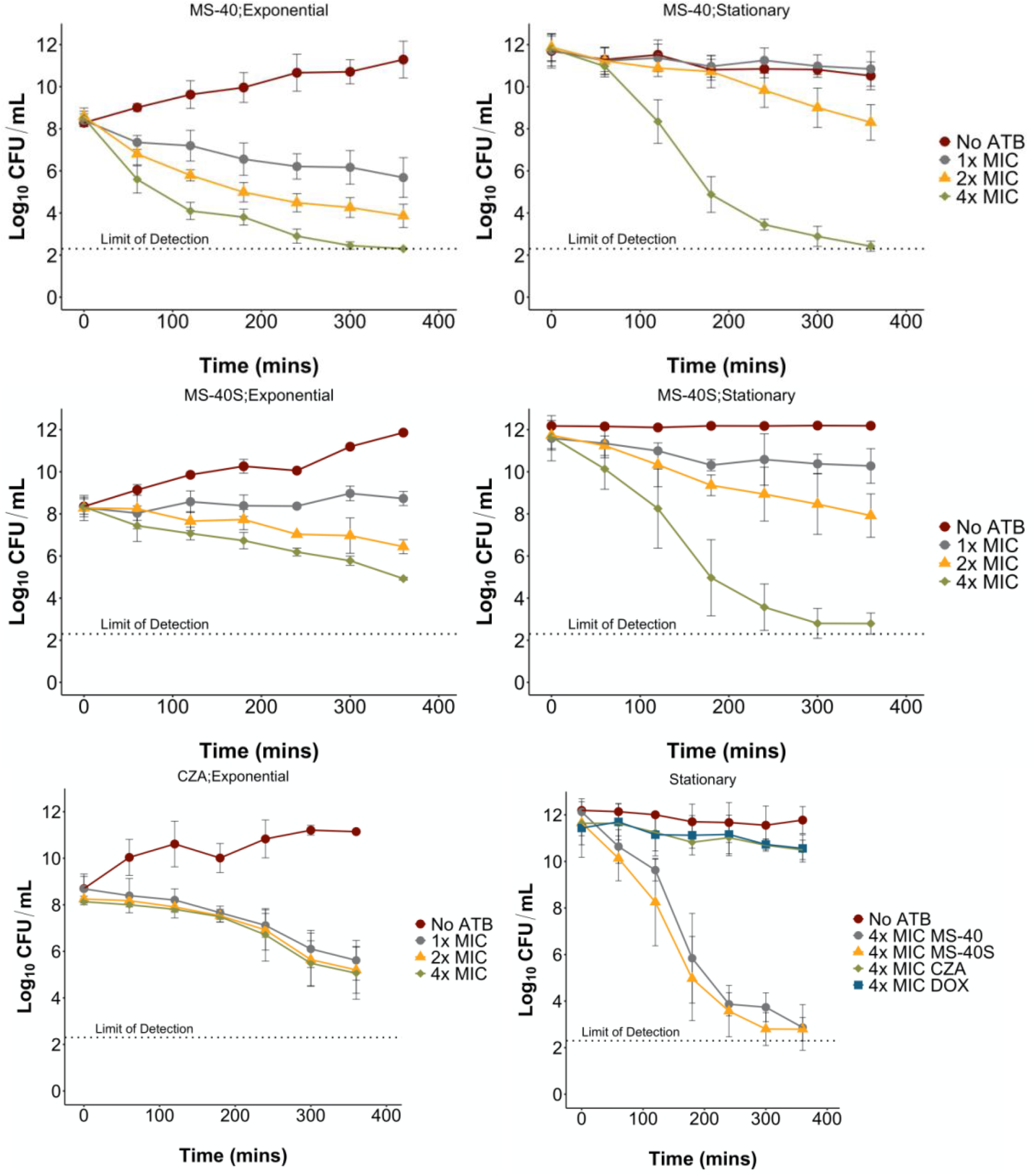
Exponential and stationary time kills of *Burkholderia cenocepacia* K56-2. The compounds used were MS-40, MS-40S, and ceftazidime-avibactam (CZA) at 1×, 2×, and 4× MIC, as well as doxycycline (DOX) at 4× MIC. The cultures were grown overnight for stationary phase cells or grown overnight,subcultured, and grown to early exponential phase. Samples were taken every hour for six hours to determine CFU/mL. No ATB; no antibiotic.

The finding that MS-40S and MS-40 is able to kill stationary phase cells highlights their potential as future therapeutics. Stationary phase cells contain a higher amount of persister cells that could be a common cause of relapses in infections (10, 34).

### MS-40S kill and inhibit the formation of persister cells

Persister cells, a subpopulation that is not killed by an antimicrobial, are thought to be a common cause of relapses in infections and persistent infections (34, 35). Persisters are thought to form via randomly overexpressing a resistance factor, decreased growth rate, decreased cellular energy, and/or a slower lag phase (35, 36). Once the antibiotic is removed they will begin to grow normally, without inherited resistance, termed “persister awakening” (36). A stationary phase population has increased amounts of persisters because they are slower growing and are metabolically dormant (34). MS-40S and MS-40 can kill stationary phase cells effectively, so we reasoned that the compounds might kill persister cells.

We exposed an exponentially growing *B. cenocepacia* K56-2 population to 5× MIC of ciprofloxacin (MIC, 2 μg/mL) for 3 hours, to enrich the surviving population in persister cells. After the treatment, surviving cells were washed and collected in phosphate buffered saline (PBS) to prevent persister awakening (37), then exposed to the MS-40S and MS-40. Figure 4 show persister cells, in the presence of MS-40S and MS-40 (Figure 4, top panels), are killed to a concentration below/close to the limit of detection, whereas the persister cells re-exposed to ciprofloxacin or those without antibiotics are not killed. This demonstrates that MS-40S and MS-40 can indeed kill persister cells created by other antibiotics.

**Figure 4.**
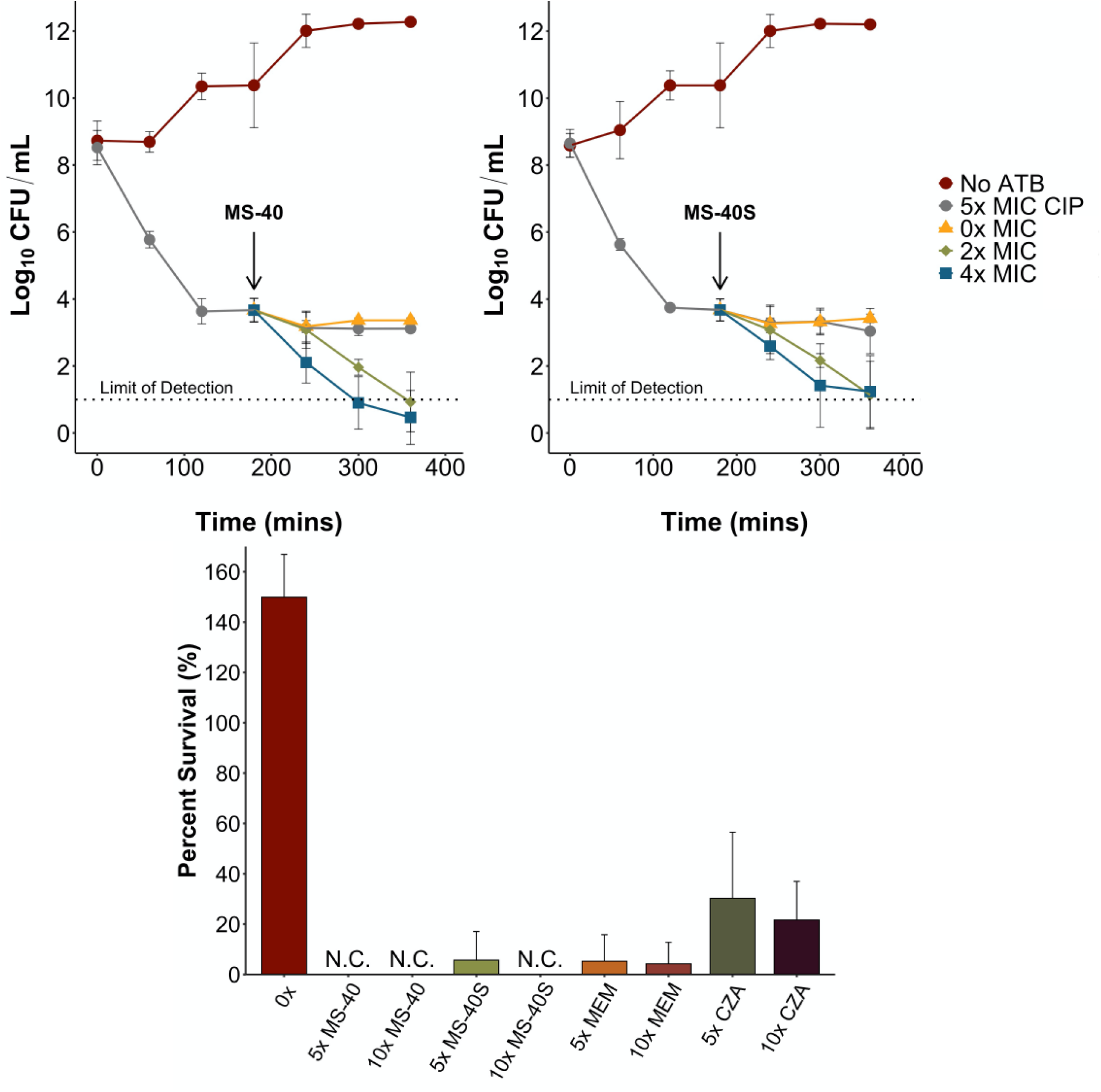
MS-40S kills and inhibits the formation of persister cells. (Top) An exponential phase culture with approximately 1×10^8^ CFU/mL were exposed to 5× MIC ciprofloxacin (CIP) to generate persister cells. At three hours post exposure, persister cells were washed, resuspended in PBS and exposed to 0×, 2×, or 4× MIC of MS-40 (top left) and MS-40S (top right) for an additional three hours. No ATB; no antibiotic. (Bottom) An overnight culture of *B. cenocepacia* K56-2 was incubated with the corresponding antimicrobial, MS-40, MS-40S, meropenem (MEM), or ceftazidime-avibactam (CZA). Percent survival was calculated by the log_10_ CFU/mL of the surviving population/log_10_ CFU/mL of the initial population. N.C., no colonies.

To determine the amount of persisters remaining after 24 hours of exposure, we performed the persister frequency assay (38). We exposed a culture with a CFU/mL of 1×10^8^, to one of the antimicrobials for 24 hours. Compounds tested included MS-40S, meropenem, ceftazidime-avibactam, and MS-40. The remaining cells, enriched in persisters, were plated on LB to determine CFU/mL and percent survival was calculated by the log_10_ CFU/mL values. After 24 hours, no persisters were formed after exposure to 10× MIC MS-40S, and a very low persister frequency with 5× MIC MS-40S (Figure 4, bottom). No persisters were formed exposed to 5× and 10× MIC of MS-40 and meropenem also produced a low amount of persisters at both concentrations tested. The last resort combination therapy ceftazidime-avibactam (CZA) produced the most persisters in this assay, with approximately 30% of the culture enriched in persisters surviving after treatment.

Taken together, these results show MS-40S, as well as MS-40, can kill persister cells created by other antibiotics and can inhibit persister cell formation. This suggests that MS-40S, as well as MS-40, have the potential to effectively eradicate an infection, reducing the risk of a relapse in infection after the treatment regime.

### *C. elegans* and *G. mellonella* Toxicity

In preliminary cytotoxicity tests, MS-40 was shown to have lower toxicity in human A549 cells than auranofin (21,22); however, the novel MS-40S has not been tested. To show that these compounds are safe for eukaryotic cells, we first used *C. elegans* as a model organism.

We performed a survival assay with *Caenorhabditis elegans* exposed to MS-40S, and three clinical antibiotics: the combination ceftazidime-avibactam, meropenem, and doxycycline, as well as MS-40 (Table 3). We calculated the Survival_100_/MIC value, which is a ratio of the highest concentration with 100% survival to the compound’s MIC (39). This is a preliminary view to a compound’s therapeutic index. MS-40S have similar Survival_100_/MIC values as clinical antibiotics, with MS-40S, doxycycline, and ceftazidime-avibactam having values of 8, 16, and 32, respectively, and MS-40 and meropenem having a value of 4. Similar to *C. elegans, Galleria* larvae were also well tolerated to MS-40S (Figure 5). This was compared to MS-40 and a clinical antibiotic, doxycycline, which has similar MIC values to MS-40S and MS-40. Concentrations used ranged from 10 to 1 mg/kg. MS-40S and MS-40 were safe for the larvae at concentrations of 10 mg/kg with a percent survival of approximately 80%. This was similar as the clinical antibiotic, doxycycline. Taken together, *C. elegans* and *Galleria* toxicity models show that MS-40S, as well as MS-40, have low toxicity in these eukaryotic organisms.

**Table 3.** Percent survival of *C. elegans* exposed to MS-40, MS-40S, and clinical antibiotics.

| Antibiotic | Concentrations (µg/mL) |  |  |  |  |  |  |  | Survival<sub>100</sub>/MIC ratio* |
| --- | --- | --- | --- | --- | --- | --- | --- | --- | --- |
|  | 256 | 128 | 64 | 32 | 16 | 8 | 4 | DMSO control |  |
| MS-40 | 74.4 | 81.8 | 94.7 | 92.9 | 100 | 100 | **100** | 100 | 4 |
| MS-40S | 85.7 | 97.6 | 100 | 100 | 100 | **100** | 100 | 100 | 8 |
| MEM | 93.2 | 97.4 | 100 | 100 | **100** | 100 | 100 | 100 | 4 |
| DOX | 79.5 | 91.5 | 100 | 98.2 | 100 | 97.4 | **100** | 100 | 16 |
| CZA<sup>a</sup> | 88.2 | 100 | 100 | 100 | 100 | 100 | **100** | 100 | 32 |

The values in bold indicate percent survival of the compound at its MIC value. \*The Survival<sub>100</sub>/MIC ratio is calculated as the highest concentration in which there is 100 percent survival divided by the compound's MIC. <sup>a</sup>Ceftazidime with 8 µg/mL avibactam. Values reported are the mean values from three independent replicates.

**Figure 5.**
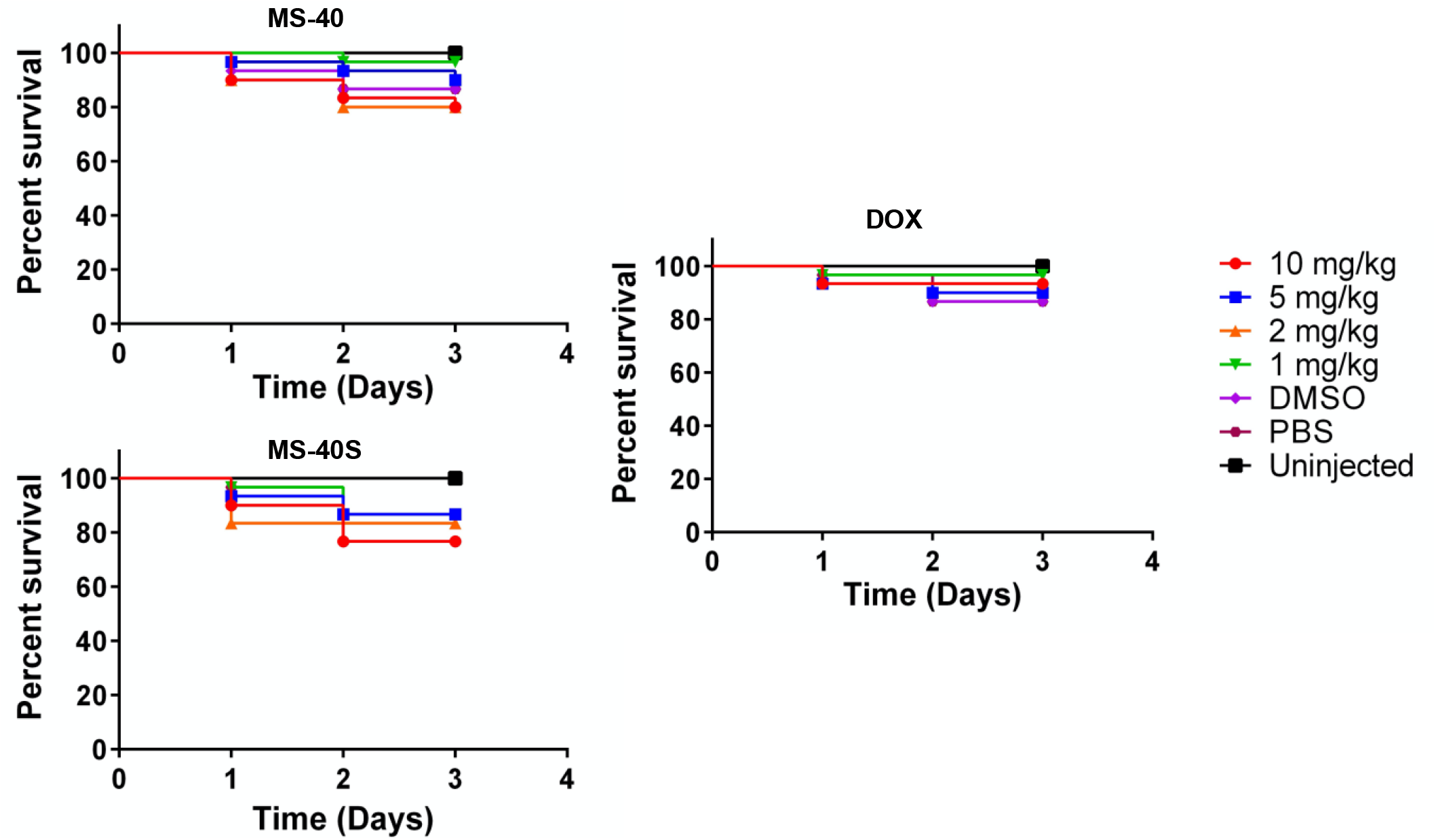
*Galleria* larvae are well tolerated to MS-40S. 10 *Galleria* larvae were injected with 10 μL of 4 concentrations of each antimicrobial, 10-1 mg/kg. 10 additional larvae were injected with 10 μL of a DMSO solvent control, PBS or uninjected. Data represents three biological replicates.

### MS-40S has Broad Bactericidal Activity

Individuals with CF are commonly infected by multiple bacteria, causing polymicrobial infections (40, 41). Therefore, for MS-40S and MS-40 to be effective antimicrobials, it is imperative for these compounds to kill additional CF pathogens. Common bacteria that cause CF lung infections, besides *Burkholderia spp*., are *Pseudomonas aeruginosa, Staphylococcus aureus, Stenotrophomonas maltophilia*, and *Achromobacter xylosoxidans* (40, 42). Other pathogenic Gram-negative bacteria that were shown to, although rarely, cause CF pulmonary infections include *Escherichia coli* (43), *Escherichia vulneris* (44), *Klebsiella pneumoniae* (42), and *Acinetobacter* species (44).

We thus tested MS-40S against CF pathogens to determine their MIC and MBC (minimum bactericidal concentration), and the same was done with MS-40 (Table 4). MS-40S have low MICs for the Gram-positive bacterium *S. aureus*, one of which is a MRSA strain. MS-40S is also bactericidal against the other Gram-negative bacteria tested with MICs in the range of 1-16 μg/mL and most of the MBCs between 1- and 4-fold their respective MICs. *P. aeruginosa*, a common multi-drug resistant bacterium (45), some strains/clinical isolates being extensively-drug resistant (46), have moderate MIC values between 16 and 64 μg/mL and their MBCs between 2- and 16-fold higher than their MICs. MS-40 shows similar results to MS-40S. Overall, the MICs/MBCs against CF pathogens show MS-40S and MS-40 have broad-spectrum bactericidal activity, indicating their potential as a therapeutic option for CF patients.

**Table 4.**
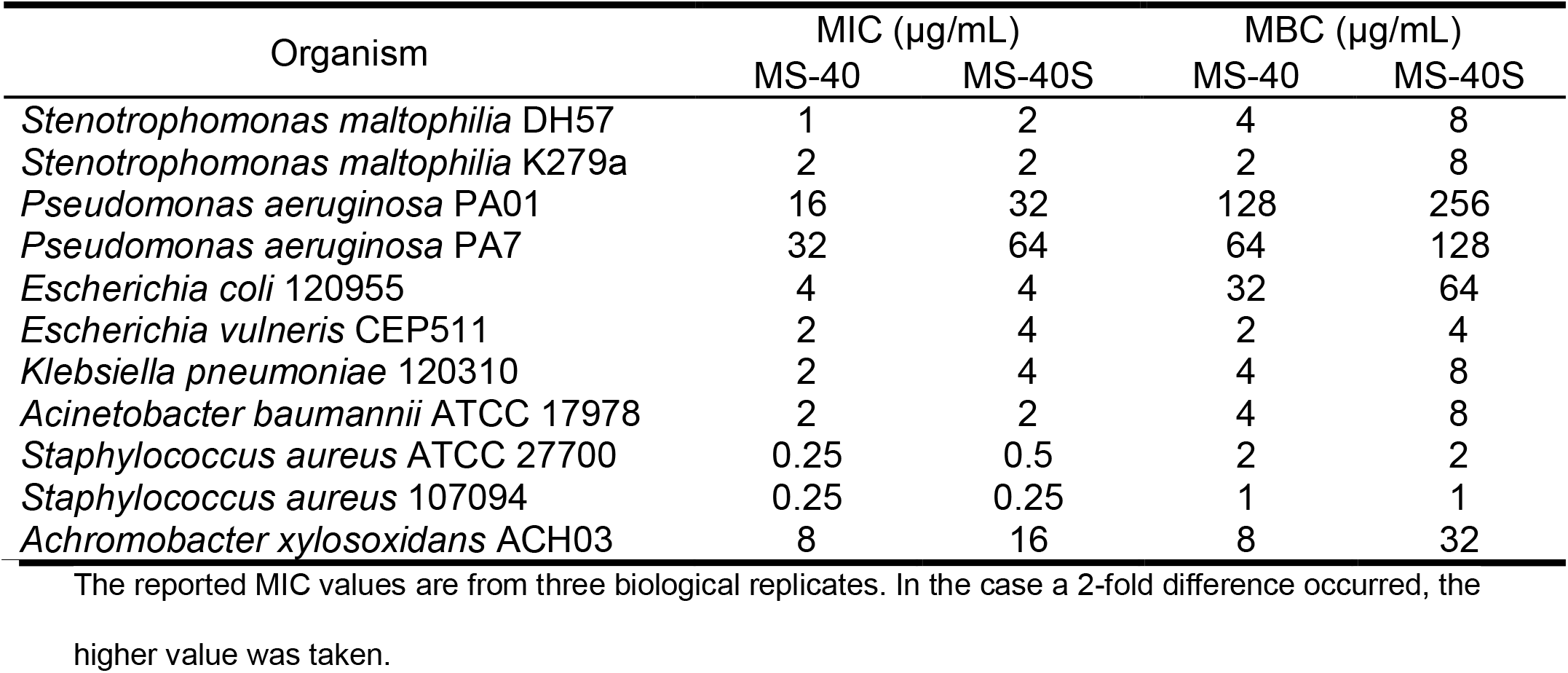
MS-40S is bactericidal against other bacteria that infect CF patients.

| Organism | MIC (µg/mL) |  | MBC (µg/mL) |  |
| --- | --- | --- | --- | --- |
|  | MS-40 | MS-40S | MS-40 | MS-40S |
| *Stenotrophomonas maltophilia* DH57 | 1 | 2 | 4 | 8 |
| *Stenotrophomonas maltophilia* K279a | 2 | 2 | 2 | 8 |
| *Pseudomonas aeruginosa* PA01 | 16 | 32 | 128 | 256 |
| *Pseudomonas aeruginosa* PA7 | 32 | 64 | 64 | 128 |
| *Escherichia coli* 120955 | 4 | 4 | 32 | 64 |
| *Escherichia vulneris* CEP511 | 2 | 4 | 2 | 4 |
| *Klebsiella pneumoniae* 120310 | 2 | 4 | 4 | 8 |
| *Acinetobacter baumannii* ATCC 17978 | 2 | 2 | 4 | 8 |
| *Staphylococcus aureus* ATCC 27700 | 0.25 | 0.5 | 2 | 2 |
| *Staphylococcus aureus* 107094 | 0.25 | 0.25 | 1 | 1 |
| *Achromobacter xylosoxidans* ACH03 | 8 | 16 | 8 | 32 |

The reported MIC values are from three biological replicates. In the case a 2-fold difference occurred, the higher value was taken.

## Discussion

Here, we show initial antibiotic properties of two auranofin analogs, MS-40 and the novel compound MS-40S, against the cystic fibrosis pathogen *B. cenocepacia* K56-2. The antibiotic properties were explored in parallel with commonly used antibiotics, namely doxycycline, meropenem, and ceftazidime-avibactam. This comparison shows MS-40S and MS-40 to have potential to be developed as antibiotics. One difference between MS-40S and MS-40 and the antibiotics used in this study is the ability of MS-40S and MS-40 to eliminate non-replicating cells. It is common for antibiotics to act on essential targets, such as involved in cell wall synthesis, DNA replication, and translation (31). In stationary phase, most of these processes are decreased, preventing the antibiotics from acting upon the cell. We confirmed this with two antibiotics with different mechanisms of action (MOA): doxycycline, a tetracycline that binds to the 30s subunit of the ribosome, preventing translation elongation (47), and ceftazidime-avibactam, a cephalosporin β-lactamase inhibitor combination that inhibits cell wall synthesis (48). These two antibiotics did not kill cells in stationary phase. Our data shows that MS-40S and MS-40 are bactericidal against both replicating and non-replicating cells. Interestingly, MS-40S was shown to kill a greater amount of stationary phase cells than exponential phase, unlike MS-40. This suggests that the MOA of these two compounds may be slightly different.

*B. cenocepacia* strains are inherently resistant to many available antibiotics (6, 10, 49), leaving only a few available for treatment. As from the resistance studies, resistance is not easy to achieve for MS-40S and MS-40. Mutational resistance can be achieved by altering the antibiotic gene targets, decreasing the binding affinity of the antimicrobial to the gene product, decrease in uptake/increase in efflux, or, lastly, by changing global responses such as changing a metabolic pathway (3). Meropenem and doxycycline quickly generated multistep resistance, possibly by one of the mechanisms listed above; however, resistance did not emerge for MS-40S and MS-40. This might be due to MS-40S and MS-40 not being affected by the change of porins/efflux pumps that can cause resistance to other antimicrobials (3), especially in *Burkholderia* species (6, 10, 50). Alternately, mutations in the gene targets of MS-40S and MS-40 could have resulted in a reduced fitness of the resistant mutant cell, preventing the mutant from outcompeting the sensitive cells (51,52).

Additionally, MS-40S and MS-40 can clear difficult to eradicate persister cells which commonly causes relapses in infections (35). This suggests these compounds could eradicate the difficult to treat persistent infections in the CF lung, helping CF patients infected with *Burkholderia* species become eligible for lung transplants by (11). MS-40S and MS-40 were shown to eliminate persister cells in two ways. First by killing an enriched population of ciprofloxacin-generated persisters, reducing the population by a further 1-2 log_10_ CFU/mL. The second way was from a stationary-phase population of bacteria exposed solely to MS-40S and MS-40, with MS-40S only having a small amount of persisters at 5× MIC and MS-40 producing no persisters. Alternatively, the persister frequency was low for meropenem (5-7%), and approximately 30% for ceftazidime-avibactam, similar to the amount produced by *Burkholderia pseudomallei* exposed to ceftazidime (38). Therefore, MS-40S and MS-40 could be used on their own to eliminate infections, or in tandem with current antibiotics to help eradicate infections (53).

We have also showed MS-40S and MS-40 have low toxicity in *C. elegans* and *G. mellonella*. One limitation of these compounds is that we did not observe *in vivo* antibiotic activity in *C. elegans* and *G. mellonella* infected with *B. cenocepacia* K56-2 (data not shown). To help explain why we were not seeing a protective effect, we tested the MIC of MS-40S and MS-40 in 50% human serum. In the presence of 50% human serum, the MIC’s of both compounds increased to 128 μg/mL, suggesting these compounds bind non-specifically to proteins, which could decrease their antimicrobial activity. To increase the efficacy of these potent antimicrobials, a drug delivery system must be developed. One possible route is the creation of a MS-40S/MS-40-loaded liposome, as has been shown with rifampicin in the treatment of pulmonary *Mycobacterium abscessus* infections (54). Creating an aerosolized antimicrobial therapy would also allow to achieve higher concentrations of the drug, and increase lung penetration, especially important for CF pulmonary infections (55).

Auranofin, and previously published auranofin analogues, inhibit thioredoxin reductase, an enzyme that plays a role in thiol-homeostasis in the cell (20, 21). However, it is unclear whether MS-40S and MS-40 share the same target with auranofin. It is assumed that the active component of auranofin is the gold atom, which binds the sulfur in the active site of thioredoxin reductase, inhibiting the formation of the critical disulfide bond, disrupting the function of the enzyme (18, 21). Auranofin was shown to have other effects on the bacterial cell, such as inhibiting DNA, protein, and cell wall synthesis. Thus, thioredoxin reductase may not be the sole antimicrobial target (17). The other potential targets of auranofin and auranofin derivatives are not known. Other factors in the mechanism of action of the antimicrobials, which can explain differences in activity among compounds with slightly different structures can be associated efflux pumps and transporters. Future research avenues could include determining the mechanism of action of MS-40S and MS-40, including the targets of the compounds, permeability factors, such as transporters and efflux pumps, and how these compounds kill persister cells and stationary phase cells.

To conclude, we have shown MS-40 and the novel compound MS-40S have potent bactericidal activity towards pathogenic *Burkholderia*, including the cystic fibrosis multi-drug resistant pathogens from the Bcc, *B. pseudomallei* and *B. mallei*. MS-40S and MS-40 kill both *B. cenocepacia* replicating and non-replicating cells, including persister cells, with little occurrence of resistance. MS-40S and MS-40 also have bactericidal activity against other pathogens involved in the CF lung microbiome. We also demonstrate in *C. elegans* and *Galleria* models, MS-40S and MS-40 were non-toxic. The novel compounds are comparable to current clinical antibiotics used to help those infected with *B. cenocepacia* and other *Burkholderia* species. We propose that MS-40S and MS-40 have unique properties as antimicrobials and studying the mechanism of action of these will help in the development of novel antibiotics to treat multi-drug resistant CF lung infections.

## Material and Methods

### Materials

All reagents and solvents were used as received from Sigma-Aldrich or Fisher Scientific unless noted. Reactions were monitored by thin layer chromatography (TLC) using TLC plates pre-coated with silica gel 60 F_254_ (Merck KGaA, Darmstadt, Germany), visualized with a handheld ultraviolet device either directly or after staining with 5% H_2_SO_4_ in ethanol. ^1^H and ^13^C nuclear magnetic resonance (NMR) spectra were recorded on a Bruker Avance Spectrospin DRX500 spectrometer, referenced either to the non-deuterated residual solvent peaks or tetramethyl silane peak (TMS, δ 0.00 ppm).^31^P NMR spectra were recorded on a Bruker Avance Spectrospin DPX200 spectrometer, using freshly prepared triphenylphosphine solution (0.1 M in CDCl_3_, δ - 6.00 ppm) as the external standard.

### Bacterial Strains and Growth Conditions

Strains used are shown in Table S2. All strains were grown in LB at 37 °C with shaking at 230 rpm. *B. ubonensis* was grown at 30 °C with shaking at 230 rpm. New Brunswick Innova40 shaking incubator was used for liquid cultures. A Barnstead Lab-Line Standing Incubator was used for LB-agar plates and 96-well plates.

### Synthesis of Auranofin and Auranofin Derivatives

Auranofin, WB-19-HL4118 (MS-40), WB-18-FI3683, WB-16-EO9899, WB-19-HG5899 and WB-19-HB2664 were synthesized following our previously published protocol (22).

#### WB-19-HL4170 (MS-40S)

To a solution of Me_3_PAuCl (150 mg, 0.486 mmol) and 1-mercapto-2-propanol (43 μL, 0.486 mmol) in MeOH (10 mL), NaOCH_3_ was added (25 wt% in methanol, 125 μL). The solution was stirred at room temperature for 2 h. The reaction mixture was then concentrated on a rotary evaporator, diluted with dichloromethane, and poured into water followed by 3 times extraction by dichloromethane. The combined organic phase was dried over Na_2_SO_4_, concentrated, passed through a PTFE syringe filter (0.2 μm), and dried in vacuum to afford the product as light beige crystals (172 mg, 97%). ^1^H NMR (500 MHz, CDCl_3_) δ 3.70 (dqd, *J* = 9.4, 6.1, 3.4 Hz, 1H), 3.47 (s, 1H), 3.09 (dd, *J* = 12.7, 3.3 Hz, 1H), 2.82 (dd, *J* = 12.7, 9.0 Hz, 1H), 1.60 (d, *J* = 10.4 Hz, 5H), 1.24 (d, *J* = 6.1 Hz, 2H), (Figure S1). ^13^C NMR (126 MHz, CDCl_3_) δ 70.41, 38.47, 21.57, 16.03 (d, *J* = 35.7 Hz), (Figure S2). ^31^P NMR (162 MHz, CDCl_3_) δ −0.31 (Figure S3).

WB-19-HL4171, WB-19-HL4172 and WB-19-HL4181 were synthesized following the same procedure as MS-40 above.

#### WB-19-HL4171

Light yellow crystals (183 mg, 99%) from Me_3_PAuCl (150 mg, 0.486 mmol) and 1-thioglycerol (36 μL, 0.486 mmol). ^1^H NMR (500 MHz, CDCl_3_) δ 3.83 − 3.60 (m, 4H), 3.10 (dd, *J* = 12.8, 4.4 Hz, 1H), 3.00 (dd, *J* = 12.8, 7.7 Hz, 1H), 2.93 (s, 1H, OH_α_), 2.32 (s, 1H, OH_β_), 1.60 (d, *J* = 10.4 Hz, 9H), (Figure S4). ^13^C NMR (126 MHz, CDCl_3_) δ 74.49, 65.58, 32.65, 16.13 (d, *J* = 35.8 Hz), (Figure S5). ^31^P NMR (162 MHz, CDCl_3_) δ −0.36, (Figure S6).

#### WB-19-HL4172

Light grey semi-solids (175 mg, 99%) from Me_3_PAuCl (150 mg, 0.486 mmol) and 3-mercapto-1-propanol (42 μL, 0.486 mmol). ^1^H NMR (500 MHz, CDCl_3_) δ 3.83 (t, *J* = 5.9 Hz, 2H), 3.19 (s, 1H), 3.04 (t, *J* = 6.8 Hz, 2H), 1.92 (p, *J* = 6.6 Hz, 2H), 1.60 (d, *J* = 10.4 Hz, 9H), (Figure S7). ^13^C NMR (126 MHz, CDCl_3_) δ 62.30, 39.20, 25.82, 16.07 (d, *J* = 35.5 Hz), (Figure S8). ^31^P NMR (162 MHz, CDCl_3_) δ −0.41, (Figure S9).

#### WB-19-HL4181

Colorless viscous solids (113 mg, quantitative) from Me_3_PAuCl (100 mg, 0.324 mmol) and cysteamine hydrochloride (37 mg, 0.324 mmol). ^1^H NMR (500 MHz, CDCl_3_) δ 3.01 (t, *J* = 6.3 Hz, 2H), 2.88 (t, *J* = 6.3 Hz, 2H), 1.99 (s, 2H), 1.59 (d, *J* = 10.4 Hz, 9H), (Figure S10). ^13^C NMR (126 MHz, CDCl_3_) δ 47.89, 32.94, 16.02 (d, *J* = 35.6 Hz), (Figure S11). ^31^P NMR (162 MHz, CDCl_3_) δ −0.15, (Figure S12).

#### WB-19-HL4121a

To a solution of (2-mercaptoethanolato)(trimethylphosphine) gold(I) (200 mg, 0.570 mmol) in 100 mL of dichloromethane, ferrocenium hexafluorophosphate (95 mg, 0.286 mmol) was added. The solution was stirred at 0 °C for 23 h. After filtration, the filtrate was concentrated and was then transferred to a 20-mL scintillation vial with a total volume of ~ 5 mL. The solution was placed in an ether vapor environment at 4 °C overnight. The yellow crystals formed were washed by diethyl ether and were further purified by preparative silica gel TLC (10:1 v/v dichloromethane/methanol) to give the product as a white solid (22 mg, 7.5%). ^1^H NMR (400 MHz, CD_2_Cl_2_) δ 1.60 (t, *J* = 4.1 Hz, 1H), (Figure S13). ^31^P NMR (162 MHz, CD_2_Cl_2_) δ 8.2, 143.8 (septet, *J*_PF_ = 710 Hz), (Figure S14).

### Auranofin Derivatives and Antibiotic Stock Solutions

Stock solutions of the derivatives were dissolved in dimethyl sulfoxide (DMSO) at a concentration of 20 mg/mL. Antibiotics were suspended at the following concentrations: tobramycin, 10 mg/mL in H_2_O (Alfa Aesar); chloramphenicol, 20 mg/mL in ethanol (Sigma); ceftazidime, 10 mg/mL in 0.1 M NaOH (Sigma); meropenem, 10 mg/mL in DMSO (Sigma); doxycycline, 25 mg/mL in H_2_O (Sigma); and ciprofloxacin, 10 mg/mL in 0.1 M HCl (Sigma).

### Antimicrobial Susceptibility Testing and Multistep Resistance to Active Derivatives

The compounds were diluted from their stock solutions to 256 μg/mL in Cation-Adjusted Mueller Hinton Broth (CAMHB) for use in experiment. Determination of MIC was followed by standards set by the Clinical Laboratory Standards Institute (CLSI) (56). 96-well plates were filled with 50 μL of CAMHB, combined with a concentration gradient of compound to be tested. Bacterial culture was diluted to a turbidity equal to MacFarland Standard 0.5, then diluted 100-fold in CAMHB. 50 μL of culture was transferred into each well. After incubation at 37 °C with no shaking for 18 hours, MIC was read visually as the lowest concentration of antibiotic that prevented growth.

To determine the rate of multistep resistance mutations from serial passaging, the assay was performed as described previously (30, 57). From the MIC plate, 30 μL from the well that had bacterial growth at the highest concentration of the antimicrobial (0.5× MIC), for each of the compounds tested, was inoculated into 2 mL of LB without compound and incubated overnight at 37 °C with shaking. These overnight cultures were then used as the culture for a second MIC test, and this was repeated 12 times for a total of 24 days of continuous growth.

### Time Kill Assays

Bacterial cultures were grown overnight and either subcultured to an OD_600_ of 0.025 or left in stationary phase. If subcultured, the bacteria were grown to early exponential phase (OD_600_ of 0.13-0.18). The bacteria were exposed to the antibiotics at 1×, 2×, and 4× the MIC, as well as no antibiotic for a negative control. Each hour from time zero to six hours, a sample of each condition was serially diluted to a dilution factor of 10^−8^, and 5 μL of each dilution was spotted onto LB agar. Plates were incubated for 24 hours at 37 °C to determine CFU/mL.

### Time Kill of Persister Cells

The generation and collection of persister cells was adapted from Bahar et al. (37). Briefly, persister cells were generated by subculturing an overnight culture of *B. cenocepacia* K56-2 in LB and grown until it reached early exponential phase (OD_600_ of 0.13-0.18). The culture was then exposed to 5x MIC of ciprofloxacin (CIP; MIC = 2 μg/mL) with 0x MIC as a control for three hours. For the initial time zero count, a sample was taken and diluted to a factor of 10^−8^ and 5 μL was spotted onto LB. After initial count, ciprofloxacin was added to the corresponding culture. A sample was taken every hour for three hours for CFU/mL counts, as mentioned above. After the third hour, the remaining population, enriched in persister cells, was collected, washed, and resuspended in phosphate buffered saline (PBS), divided into five tubes, and again exposed to 5× MIC ciprofloxacin or different concentrations of MS-40S or MS-40 (2× and 4× the MIC), along with a no antibiotic condition as a control. Samples were taken every hour for an additional three hours. Plates were incubated at 37 °C for 24 hours and counted for CFU/mL.

### Persister Frequency Assay

The persister frequency assay was performed as described in Ross et al (38). An overnight culture of *B. cenocepacia* K56-2 was subcultured to a concentration of 1×10^8^ CFU/mL in 2 mL LB. Antimicrobials tested, meropenem, ceftazidime-avibactam, MS-40S, and MS-40 were added to a final concentration of 5× and 10× MIC. The cultures were exposed to the antibiotics for 24 hours at 37°C with shaking. After 24 hours the culture was plated on LB to determine CFU/mL. Plates were incubated at 37°C for 24 hours.

### *C. elegans* Survival

*Caenorhabditis elegans* was used as a model organism to test the toxicity of the compounds. The survival was performed as described in Selin et al (39). *C. elegans* DH26 eggs were incubated at 26°C until they reached L4 stage, approximately 48 hours. L4 stage worms were collected and washed with M9 media. Worms were suspended in 100 μL of M9 and transferred to the NGMII plates containing *E. coli* OP50. Approximately ten non-infected *C. elegans* OP50 fed worms, in triplicate, were exposed to a serial dilution of antibiotics to be tested in liquid killing media (LKM; 80% M9 buffer 20% liquid NGMII) in a 96-well plate. The range of concentrations used was 128 – 4 μg/mL for the following antimicrobials: MS-40S, MS-40, meropenem, doxycycline, and ceftazidime-avibactam, along with a no antibiotic control. Worms were counted at day 0 and incubated at 25°C. After 24 hours, worms were counted for percent survival and the Survival_100_/MIC ratio was calculated. Worms that appeared straight were considered dead, and those were moving and S-shaped were counted as alive. Three biological replicates were performed.

### *Galleria* Toxicity

*Galleria mellonella* was also used as a model organism to study the toxicity of MS-40S and MS-40. The experiments were performed as done in Naguib & Valvano 2018 (58) and Cruz et al., 2018 (59). *Galleria* larvae were stored at 16°C in wood shavings and used within 2 weeks of receipt. Larvae, with an approximate weight of 250 mg, were injected with 10 μL in the last, left proleg using a Hamilton micro-syringe (Hamilton, Nevada, USA). For each compound, MS-40S, MS-40, and doxycycline were diluted in PBS, and 10, 5, 2, and 1 mg/kg were injected in 10 worms for each condition. 10 worms were also injected with 10 μL of PBS for a negative control, as well as 10 worms were not injected. Survival was measured every 24 hours for 72 hours. Larvae were considered dead if non-motile and unresponsive to touch. Three biological replicates were performed. Survival curves were made on GraphPad Prism 6.

## Acknowledgements

Work was supported by a research grant from Cystic Fibrosis Canada and a project grant from CIHR to S.T.C., and in part by a grant from the National Institute of Allergy and Infectious Diseases (R21AI140418 to M.Y.). D. M. was supported by a scholarship from the Canadian Institutes of Health Research (CIHR). A.M.H. was supported by the Vanier Canada Graduate Scholarship. Z.L.Y. was supported by scholarship from Research Manitoba.

We would like to thank Catherine Deschênes (Laboratoire Régional de Microbiologie) for some of the *Burkholderia* isolates used in this study, to Ayush Kumar (University of Manitoba) for providing *P. aeruginosa* PA01 and PA7 strains and *A. baumannii* ATCC 17978. We would also like to thank the NMR facility in the Department of Chemistry at the University of Manitoba and ATCC for performing the MIC experiments with *B. mallei* and *B. pseudomallei*.

## Author contributions

S.T.C. conceived the idea of the research, designed the research, and edited the manuscript. M. Y. and B.W. conceived the idea of the auranofin derivatives, designed the synthesis, and edited the manuscript. D.M. designed and performed the experiments, analyzed the data, and wrote the paper. B.W., D.T., S.H.L. synthesized auranofin and the auranofin derivatives. A.M.H. aided in the experimental protocols and processing of the data. Z.L.Y aided in the design and experimental procedure of the *C. elegans* and *Galleria* toxicity. S.T.C and M.Y. supervised the work and contributed financially to the research.

## References

1. Gould K. 2016. Antibiotics: from prehistory to the present day. J Antimicrob Chemother 71:572–575.

2. Nicolaou KC, Rigol S. 2018. A brief history of antibiotics and select advances in their synthesis. J Antibiot (Tokyo) 71:153–184.

3. Munita JM, Arias CA. 2016. Mechanisms of Antibiotic Resistance, p. 481–511. In Kudva, Cornick, Plummer, Zhang, Nicholson, Bannantine, Bellaire (eds.), Virulence Mechanisms of Bacterial Pathogens, Fifth Edition. American Society of Microbiology.

4. Hutchings MI, Truman AW, Wilkinson B. 2019. Antibiotics: past, present and future. Curr Opin Microbiol 51:72–80.

5. Eberl L, Vandamme P. 2016. Members of the genus *Burkholderia*: good and bad guys. F1000Research 5:1007.

6. Rhodes KA, Schweizer HP. 2016. Antibiotic resistance in *Burkholderia* species. Drug Resist Updat 28:82–90.

7. Sfeir MM. 2018. *Burkholderia cepacia* complex infections: More complex than the bacterium name suggest. J Infect 77:166–170.

8. Lobo LJ, Noone PG. 2014. Respiratory infections in patients with cystic fibrosis undergoing lung transplantation. Lancet Respir Med 2:73–82.

9. Los-Arcos I, Len O, Martín-Gómez MT, González-López JJ, Saéz-Giménez B, Deu M, Nuvials X, Ferrer R, Román A, Gavaldà J. 2019. Lung transplantation in two cystic fibrosis patients infected with previously pandrug-resistant Burkholderia cepacia complex treated with ceftazidime–avibactam. Infection 47:289–292.

10. Scoffone VC, Chiarelli LR, Trespidi G, Mentasti M, Riccardi G, Buroni S. 2017. *Burkholderia cenocepacia* Infections in Cystic Fibrosis Patients: Drug Resistance and Therapeutic Approaches. Front Microbiol 8:1592.

11. Dupont L. 2017. Lung transplantation in cystic fibrosis patients with difficult to treat lung infections: Curr Opin Pulm Med 23:574–579.

12. Frei A, Zuegg J, Elliott AG, Baker M, Braese S, Brown C, Chen F, G. Dowson C, Dujardin G, Jung N, King AP, Mansour AM, Massi M, Moat J, Mohamed HA, Renfrew AK, Rutledge PJ, Sadler PJ, Todd MH, Willans CE, Wilson JJ, Cooper MA, Blaskovich MAT. 2020. Metal complexes as a promising source for new antibiotics. Chem Sci 11:2627–2639.

13. Chakraborty P, Oosterhuis D, Bonsignore R, Casini A, Olinga P, Scheffers D. 2021. An Organogold Compound as Potential Antimicrobial Agent against Drug-Resistant Bacteria: Initial Mechanistic Insights. ChemMedChem cmdc.202100342.

14. May HC, Yu J-J, Guentzel MN, Chambers JP, Cap AP, Arulanandam BP. 2018. Repurposing Auranofin, Ebselen, and PX-12 as Antimicrobial Agents Targeting the Thioredoxin System. Front Microbiol 9:336.

15. Novelli F, Recine M, Sparatore F, Juliano C. 1999. Gold(I) complexes as antimicrobial agents. Il Farm 54:232–236.

16. Owings JP, McNair NN, Mui YF, Gustafsson TN, Holmgren A, Contel M, Goldberg JB, Mead JR. 2016. Auranofin and N-heterocyclic carbene gold-analogs are potent inhibitors of the bacteria *Helicobacter pylori*. FEMS Microbiol Lett 363:fnw148.

17. Thangamani S, Mohammad H, Abushahba MFN, Sobreira TJP, Hedrick VE, Paul LN, Seleem MN. 2016. Antibacterial activity and mechanism of action of auranofin against multi-drug resistant bacterial pathogens. Sci Rep 6:22571.

18. Kean WF, Hart L, Buchanan WW. 1997. Auranofin. Br J Rheumatol 36:560–572.

19. Finkelstein AE, Walz DT, Batista V, Mizraji M, Roisman F, Misher A. 1976. Auranofin. New oral gold compound for treatment of rheumatoid arthritis. Ann Rheum Dis 35:251–257.

20. Harbut MB, Vilchèze C, Luo X, Hensler ME, Guo H, Yang B, Chatterjee AK, Nizet V, Jacobs WR, Schultz PG, Wang F. 2015. Auranofin exerts broad-spectrum bactericidal activities by targeting thiol-redox homeostasis. Proc Natl Acad Sci U S A 112:4453–4458.

21. Epstein TD, Wu B, Moulton KD, Yan M, Dube DH. 2019. Sugar-Modified Analogs of Auranofin Are Potent Inhibitors of the Gastric Pathogen *Helicobacter pylori*. ACS Infect Dis 5:1682–1687.

22. Wu B, Yang X, Yan M. 2019. Synthesis and Structure–Activity Relationship Study of Antimicrobial Auranofin against ESKAPE Pathogens. J Med Chem 62:7751–7768.

23. Balwan A, Nicolau DP, Wungwattana M, Zuckerman JB, Waters V. 2016. Clinafloxacin for Treatment of Burkholderia cenocepacia Infection in a Cystic Fibrosis Patient. Antimicrob Agents Chemother 60:1–5.

24. Kitt H, Lenney W, Gilchrist FJ. 2016. Two case reports of the successful eradication of new isolates of Burkholderia cepacia complex in children with cystic fibrosis. BMC Pharmacol Toxicol 17:14.

25. El-Laboudi AH, Etherington C, Whitaker P, Clifton IJ, Conway SP, Denton M, Peckham DG. 2009. Acute Burkholderia cenocepacia pyomyositis in a patient with cystic fibrosis. J Cyst Fibros 8:273–275.

26. Gilchrist FJ, Webb AK, Bright-Thomas RJ, Jones AM. 2012. Successful treatment of cepacia syndrome with a combination of intravenous cyclosporin, antibiotics and oral corticosteroids. J Cyst Fibros Off J Eur Cyst Fibros Soc 11:458–460.

27. Salizzoni S, Pilewski J, Toyoda Y. Lung Transplant for a Patient With Cystic Fibrosis and Active *Burkholderia cenocepacia* Pneumonia. Exp Clin Transplant 12.

28. Sabtu N, Enoch DA, Brown NM. 2015. Antibiotic resistance: what, why, where, when and how? Br Med Bull 116:105–113.

29. Waglechner N, Wright GD. 2017. Antibiotic resistance: it’s bad, but why isn’t it worse? BMC Biol 15:84.

30. AbdelKhalek A, Abutaleb NS, Elmagarmid KA, Seleem MN. 2018. Repurposing auranofin as an intestinal decolonizing agent for vancomycin-resistant enterococci. Sci Rep 8:8353.

31. Kohanski MA, Dwyer DJ, Collins JJ. 2010. How antibiotics kill bacteria: from targets to networks. Nat Rev Microbiol 8:423–435.

32. Kohanski MA, Dwyer DJ, Hayete B, Lawrence CA, Collins JJ. 2007. A Common Mechanism of Cellular Death Induced by Bactericidal Antibiotics. Cell 130:797–810.

33. Corona F, Martinez J. 2013. Phenotypic Resistance to Antibiotics. Antibiotics 2:237–255.

34. Wood TK, Knabel SJ, Kwan BW. 2013. Bacterial Persister Cell Formation and Dormancy. Appl Environ Microbiol 79:7116–7121.

35. Brauner A, Fridman O, Gefen O, Balaban NQ. 2016. Distinguishing between resistance, tolerance and persistence to antibiotic treatment. Nat Rev Microbiol 14:320–330.

36. Wilmaerts D, Windels EM, Verstraeten N, Michiels J. 2019. General Mechanisms Leading to Persister Formation and Awakening. Trends Genet 35:401–411.

37. Bahar AA, Liu Z, Totsingan F, Buitrago C, Kallenbach N, Ren D. 2015. Synthetic dendrimeric peptide active against biofilm and persister cells of Pseudomonas aeruginosa. Appl Microbiol Biotechnol 99:8125–8135.

38. Ross BN, Myers JN, Muruato LA, Tapia D, Torres AG. 2018. Evaluating New Compounds to Treat *Burkholderia pseudomallei* Infections. Front Cell Infect Microbiol 8:210.

39. Selin C, Stietz MS, Blanchard JE, Gehrke SS, Bernard S, Hall DG, Brown ED, Cardona ST. 2015. A Pipeline for Screening Small Molecules with Growth Inhibitory Activity against Burkholderia cenocepacia. PLOS ONE 10:e0128587.

40. Huang YJ, LiPuma JJ. 2016. The Microbiome in Cystic Fibrosis. Clin Chest Med 37:59–67.

41. Cribbs SK, Beck JM. 2017. Microbiome in the pathogenesis of cystic fibrosis and lung transplant-related disease. Transl Res 179:84–96.

42. Vandeplassche E, Tavernier S, Coenye T, Crabbé A. 2019. Influence of the lung microbiome on antibiotic susceptibility of cystic fibrosis pathogens. Eur Respir Rev 28:190041.

43. Edwards BD, Somayaji R, Greysson-Wong J, Izydorczyk C, Waddell B, Storey DG, Rabin HR, Surette MG, Parkins MD. 2020. Clinical Outcomes Associated With Escherichia coli Infections in Adults With Cystic Fibrosis: A Cohort Study. Open Forum Infect Dis 7:ofz476.

44. Burns JL, Emerson J, Stapp JR, Yim DL, Krzewinski J, Louden L, Ramsey BW, Clausen CR. 1998. Microbiology of Sputum from Patients at Cystic Fibrosis Centers in the United States. Clin Infect Dis 27:158–163.

45. Pang Z, Raudonis R, Glick BR, Lin T-J, Cheng Z. 2019. Antibiotic resistance in Pseudomonas aeruginosa: mechanisms and alternative therapeutic strategies. Biotechnol Adv 37:177–192.

46. Horcajada JP, Montero M, Oliver A, Sorlí L, Luque S, Gómez-Zorrilla S, Benito N, Grau S. 2019. Epidemiology and Treatment of Multidrug-Resistant and Extensively Drug-Resistant *Pseudomonas aeruginosa* Infections. Clin Microbiol Rev 32:e00031–19, /cmr/32/4/CMR.00031-19.atom.

47. Grossman TH. 2016. Tetracycline Antibiotics and Resistance. Cold Spring Harb Perspect Med 6:a025387.

48. Shirley M. 2018. Ceftazidime-Avibactam: A Review in the Treatment of Serious Gram-Negative Bacterial Infections. Drugs 78:675–692.

49. Ostermann MF, Neubauer H, Frickmann H, Hagen RM. 2016. Correlation of *rpsU* gene sequence clusters and biochemical properties, GC—MS spectra and resistance profiles of clinical *Burkholderia* spp. isolates. Eur J Microbiol Immunol 6:25–39.

50. Guglierame P, Pasca MR, Rossi ED, Buroni S, Arrigo P, Manina G, Riccardi G. 2006. Efflux pump genes of the resistance-nodulation-division family in *Burkholderia cenocepacia* genome. BMC Microbiol 6:66.

51. Hughes D, Andersson DI. 2017. Evolutionary Trajectories to Antibiotic Resistance. Annu Rev Microbiol 71:579–596.

52. Durão P, Balbontín R, Gordo I. 2018. Evolutionary Mechanisms Shaping the Maintenance of Antibiotic Resistance. Trends Microbiol 26:677–691.

53. Zheng EJ, Stokes JM, Collins JJ. 2020. Eradicating Bacterial Persisters with Combinations of Strongly and Weakly Metabolism-Dependent Antibiotics. Cell Chem Biol S2451945620303378.

54. Rinaldi F, Hanieh PN, Sennato S, Santis FD, Forte J, Fraziano M, Casciardi S, Marianecci C, Bordi F, Carafa M. 2021. Rifampicin–Liposomes for Mycobacterium abscessus Infection Treatment: Intracellular Uptake and Antibacterial Activity Evaluation 13.

55. Chalmers JD, van Ingen J, van der Laan R, Herrmann J-L. 2021. Liposomal drug delivery to manage nontuberculous mycobacterial pulmonary disease and other chronic lung infections. Eur Respir Rev 30:210010.

56. 2015. Methods for dilution antimicrobial susceptibility tests for bacteria that grow aerobically: M07-A10; approved standard10. ed. Committee for Clinical Laboratory Standards, Wayne, PA.

57. Yarlagadda V, Akkapeddi P, Manjunath GB, Haldar J. 2014. Membrane Active Vancomycin Analogues: A Strategy to Combat Bacterial Resistance. J Med Chem 57:4558–4568.

58. Naguib MM, Valvano MA. 2018. Vitamin E Increases Antimicrobial Sensitivity by Inhibiting Bacterial Lipocalin Antibiotic Binding. mSphere 3:e00564–18, /msphere/3/6/mSphere564-18.atom.

59. Cruz L, Lopes L, de Camargo Ribeiro F, de Sá N, Lino C, Tharmalingam N, de Oliveira R, Rosa C, Mylonakis E, Fuchs B, Johann S. 2018. Anti-Candida albicans Activity of Thiazolylhydrazone Derivatives in Invertebrate and Murine Models. J Fungi 4:134.

